# Towards a robust comparison of diversity between sampled TCR repertoires

**DOI:** 10.1101/2023.02.10.528010

**Authors:** Peter C. de Greef, Rob J. de Boer

## Abstract

T-cell receptor (TCR) repertoire sequencing data provides quantitative insight into the distribution of T-cell clones. The diversity of the TCR repertoire in humans tends do decrease with age, which may be a key determinant explaining immune senescence in older individuals. To address this, we first analyze how the diversity of a potential T-cell response against an unseen pathogen changes with age. Next, we discuss the complications with interpreting the outcomes of such an analysis. Specifically, the changes in T-cell subset sizes confound analyses of TCR diversity, and typical sample sizes do not easily allow for a robust quantification of this diversity. Thus, explaining immune senescence as a result of decreasing TCR diversity is far from straightforward and requires a detailed, robust, and quantitative analysis.

## Introduction

The human TCR repertoire has a unique composition that results from thymic production and selection, as well as exposure to antigens in the periphery. The extreme diversity of TCR sequences makes comparison of the TCR repertoire between individuals a challenging task. In addition, the Human Leukocyte Antigens (HLAs) are highly polymorphic in the human population, implying that a TCR may bind different antigens in other individuals. A measure that summarizes the entirety of a TCR repertoire is its diversity. In ecological studies, diversity is often measured considering the richness, which is defined as the total number of species in a system, and the evenness, which quantifies to which extent these species differ among each other in frequency [3]. In a TCR repertoire context, the species are the distinct TCR sequences, and their diversity can be estimated using high-throughput TCR sequencing.

The generation of new T-cell clones decreases with age, to an extent that naive T-cell production by the thymus is diminished [13] or even absent [11] in older individuals. This decreasing source of new diversity may lead to reduced T-cell immunity since TCR diversity is a key feature of a functional T-cell pool. The scenario that the TCR repertoire lacks T-cells that are specific for a given foreign antigen has been described as ‘holes in the repertoire’ [14]. Such an absence of required T-cell specificities may be the result of reduced TCR richness, illustrating the need for accurate estimates of this richness. Here we present an intuitive analysis in which we estimate how many T-cell clones would be recruited against an unseen pathogen. This case study provides quantitative insights into the potential losses of responses with age, but also highlights key caveats of such an analysis. We discuss important challenges of estimating TCR diversity based on sequencing data of sampled T cells. These insights will help to refine future experiments and analyses to better compare the TCR diversity between sampled repertoires.

## Results and Discussion

### The estimated richness of a putative response and the total repertoire decreases with age

A reduced TCR repertoire diversity, leading to a failure to mount a T-cell response, should be reflected in a strongly reduced number of TCR sequences specific for a given pathogen. We tested this by using the VDJdb, which is a database that lists TCR sequences that are found to be specific for certain epitopes [10]. We checked the occurrence of such sequences specific for a particular pathogen across a large cohort of individuals up to an age of about 70 years [6]. Since the vast majority of the population in Western countries is HIV-negative, HIV-1 will be an unseen pathogen to most individuals in this dataset. So, the number of TCRβ sequences specific for an HIV-1 epitope serves as a proxy for a putative T-cell response against a pathogen without previous exposure. This is important, as both the VDJdb and the TCR repertoires in this dataset will be enriched for specificities towards common pathogens. We counted the number of matches between the HIV-1 entries in the VDJdb and the TCRβ repertoires that were size-normalized to exclude heterogeneous sample sizes as a confounding factor (see Methods). Interestingly, although the data only covers a tiny portion of each total T-cell repertoire, we found such sequences in all individuals, across the entire age range (Fig. 1A).

**Figure 1:**
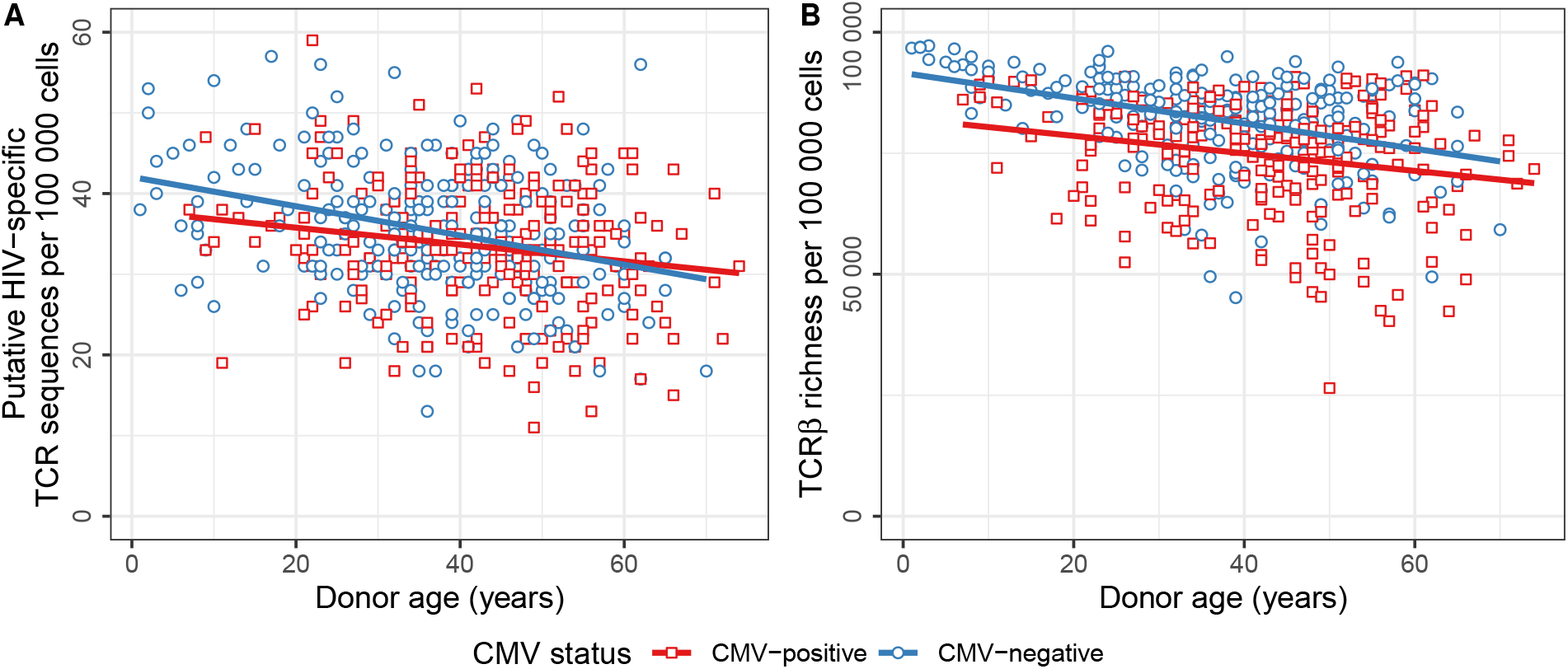
The richness of unsorted T-cell repertoires tends to decrease with age and CMV. **A**. Total number of distinct TCRβ nucleotide sequences from [6] that match the VDJdb [10] as being specific for an HIV-1 epitope in CMV-positive (red boxes) and CMV-negative (blue circles) individuals. TCRβ sequences are counted as a match with the database if their CDR3 amino acid sequence as well as their V- and J-gene families are identical. Differences in sample size are normalized for by randomly sampling a 100 000 templates from each complete sample (see Methods). **B**. Total number of distinct TCRβ nucleotide sequences among 100 000 unsorted T cells. Linear regression lines are shown for CMV-positive (red) and CMV-negative (blue) individuals.

The richness of the putative HIV-1 response varies considerably between donors, even of the same age. This reflects a large individual heterogeneity, for example due to different HLA compositions within the cohort. A linear regression analysis, with age as the independent variable, showed that the number of putative HIV-1-responsive clones tends to decline, both with age and CMV-infection (Fig. 1A). This implies that the number of responding clones tends to decrease with age, and that at an age of 60 years about a quarter of the diversity found in children is lost. The data also suggest that CMV-infection reduces the number of HIV-1-responsive clones, probably due to the repertoire being skewed towards a limited number of expanded CMV-specific T-cell clones. Note that the TCRα chain was not sequenced but also determines the TCR specificity, and that being reported as binding to epitopes may be restricted to HLA alleles that are absent in many donors. Our analysis thus does not exactly quantify the T-cell response against HIV-1 epitopes. However, the decrease in reported TCR hits is expected to reflect an actual decrease in richness of the T-cell response against an unseen pathogen. To place these pathogen-specific results into a more general context, we also quantified the overall changes in repertoire richness with age. In line with previous studies [1, 8, 9, 15], we found a moderate decline in richness per normalized number of sequences with both age and CMV-infection (Fig. 1B).

### TCR repertoires are dominated by naive T cells in young and by memory T cells in older individuals

Importantly, the TCRβ dataset we used consists of sequences that were observed in a peripheral blood sample, that was not sorted to only contain a specific T-cell subpopulation. Naive T-cell frequencies are a major determinant of TCR repertoire richness because encounter with antigen leads to proliferation, and the resulting effector and/or memory clones will mostly persist at a higher frequency than the initial frequency of the naive T-cell clone. It is important to make a distinction between the number of naive T cells per unit of blood, and the percentage of naive T cells among other subsets. For example, when the memory compartment grows with age, the naive T-cell diversity does not have to decrease if its total pool size remains stable, while the percentage of naive T cells would decrease. In line with this, Wertheimer et al. reported that the percentage of naive T-cells decreases significantly with age, while the absolute number of naive CD4 T cells per µl of blood is rather similar between young and older individuals, if they are CMV-negative [12].

We consulted various studies reporting the number of naive T cells per volume of blood to estimate how the naive T-cell pool size changes with age (Fig. 2A). The reported counts vary widely between individuals and different studies. Part of this variation may be explained by the different markers that are used to sort the naive sub-population from blood, some being more stringent than others. The studies mostly agree on the observation that naive CD8 T-cell numbers in blood decrease strongly with age (Fig. 2A; right), implying that the total number of naive CD8 T cells is much smaller in older than in young individuals. However, some of the naive CD8 T cells may have undergone phenotypic changes without having responded to foreign antigen, for example becoming virtual memory cells. The effects of age on the naive CD4 T-cell numbers in blood are less pronounced (Fig. 2A; left), suggesting that the absolute size of the naive CD4 T-cell pool does not change dramatically with age, while the relative frequency of naive CD4 T cells may decrease substantially.

**Figure 2:**
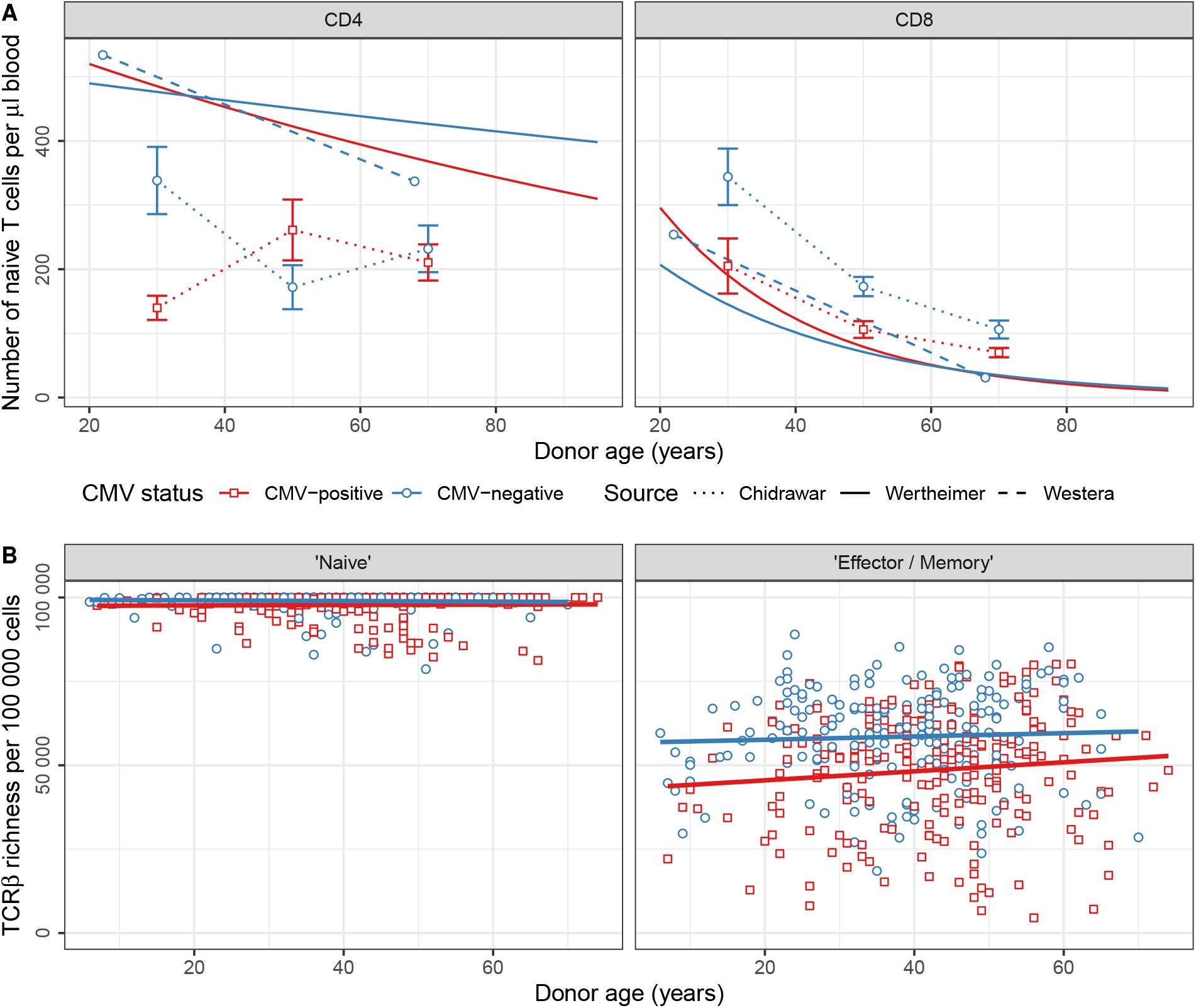
The observed T-cell diversity largely depends on the relative frequency of naive T cells. **A**. Estimated number of naive CD4 and CD8 T cells per µl of blood as reported in three studies. Solid lines show the results of a regression analysis [12], the dashed line indicates the median values of a young and aged age group [13], and the dotted lines connect the median values per age group, with the error bars indicating the reported standard deviation [4]. **B**. Size-normalized richness (similar to Fig. 1B) after splitting the TCRβ repertoires into a ‘naive’ and ‘effector/memory’ fraction (see Methods). Linear regression lines are shown for CMV-positive (red) and CMV-negative (blue) individuals.

To assess to which extent the observed changes in richness as described above may reflect relative changes in subset sizes, we defined a simple subset classifier. We split each TCR repertoire into a putative naive and effector/memory fraction (see Methods). We again plotted the size-normalized richness of each repertoire, like in Fig. 1B, but now separately for the two inferred subpopulations (Fig. 2B). The absence of a decrease in TCR repertoire richness in both compartments suggests that the diversity of each subpopulation may be rather stable with age. Although the separation between naive and effector/memory will be far from perfect with this approach, it reveals a key determinant of any TCR diversity analysis. By a relative increase of expanded subpopulations, the richness of an unsorted repertoire will decrease, while the richness of the individual subpopulations can remain stable. This means that it is crucial to analyze such subpopulations separately to allow for conclusions on diversity loss within for example the naive T-cell compartment.

### The total richness of the naive T cell repertoire can only be estimated by combining multiple subsamples

To analyze the changes in naive TCR repertoire richness with age we re-analyzed the TCRβ sequencing data analyzed before in [9]. They estimated the richness of naive and memory populations from four young (20-35y) and five aged (70-85y) individuals. Importantly, the cells were split into multiple subsamples before mRNA extraction, enabling the use of the Chao2 estimator to impute the richness of each total T-cell pool. They estimated naive TCRβ repertoire richness in the order of 10^7^ to 10^8^, with a two-to fivefold richness decrease in older healthy donors when compared to the young individuals [9]. We revisited this naive T-cell data by performing additional analyses on the changes with age. First, we estimated the relative contribution of sequence reads by single cells and used this to impute the distribution of cells from the read counts in each sample (see Methods). We then plotted the measured and extrapolated richness using rarefaction curves to account for differences in sampling depth (Fig. 3). In line with the results in Fig. 2B, the TCR richness of a given number of naive T cells is very similar between the age groups, especially for lower numbers of cells (Fig. 3A). At a depth of 50 000 cells, which was reached in all subsamples, the maximum richness difference between any sample pair was only 13% for naive CD8 T cells, and less than 5% for CD4. Although the curves diverge somewhat more at a higher depth, the observed differences in measured richness between the age groups remain rather limited.

**Figure 3:**
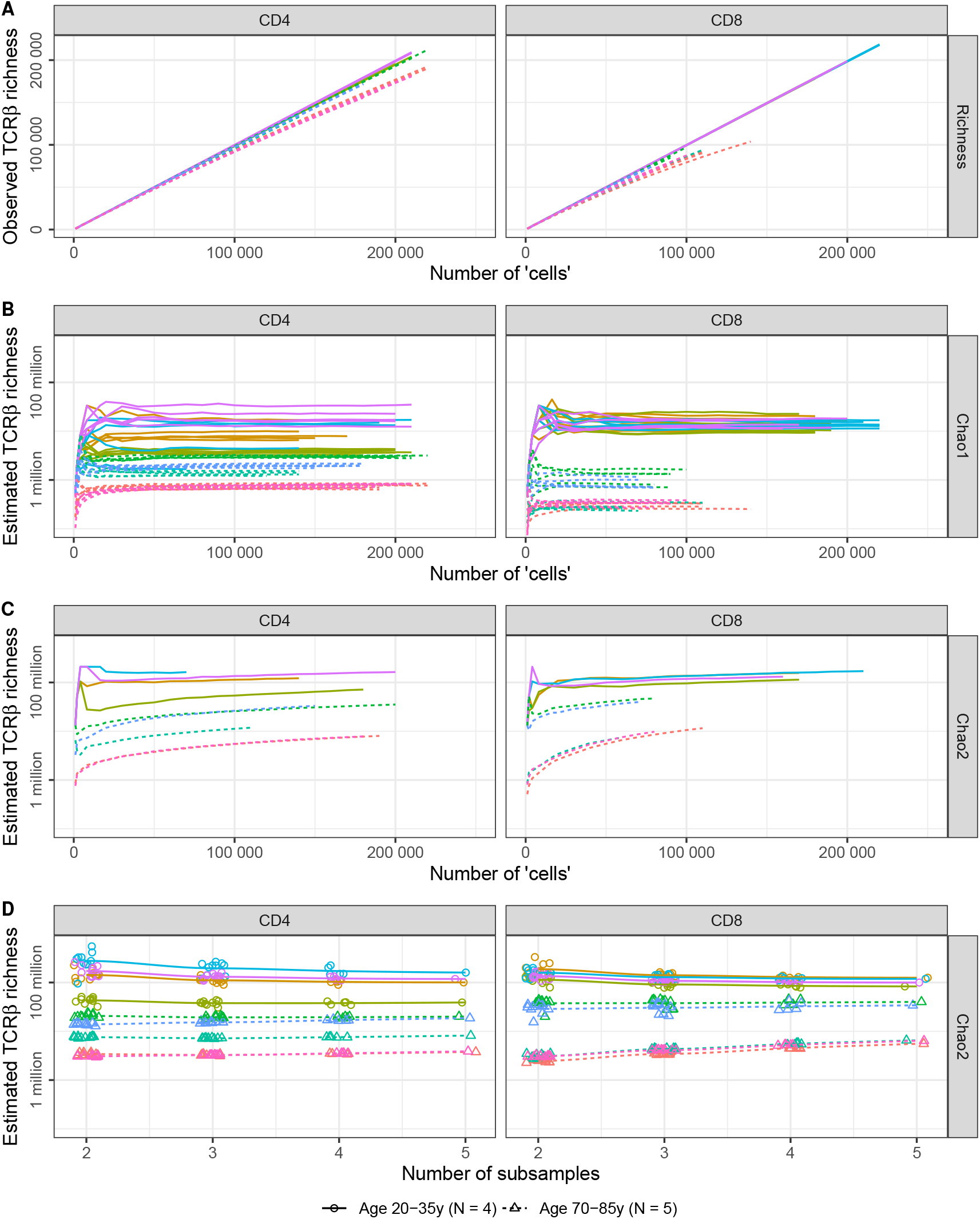
Naive TCRβ richness in young and aged individuals as a function of sample size and number. **A-C**. Rarefaction curves for the observed TCRβ richness (A) and estimated total richness using the Chao1 (B) and Chao2 (C) estimators. Shown are results based on CD4 (left) and CD8 (right) naive T-cell repertoires from young (solid lines) and aged individuals (dashed lines). The colors indicate individual donors, and the horizontal axis depicts the inferred number of cells (see Methods). The rarefaction curve of the Chao2 estimator runs until the sampling depth of the smallest subsample of each individual. **D**. Estimated total TCRβ richness using the Chao2 estimator based on 50 000 inferred ‘cells’ from all (5) or a subset of the subsamples. Shown are the loess regression lines with colors and line style as in A-C, with the symbols showing individual Chao2 estimates for repertoires of young (circles) and aged (triangles) individuals.

Since typical samples of T cells only comprise a minor fraction of the entire T-cell repertoire, it can be useful to extrapolate the richness to the level of the entire T-cell pool. The non-parametric Chao1 estimator accounts for the unobserved diversity, based on the number of TCR sequences observed once and twice in a sample [2].

For each subsample, the total richness estimate using the Chao1 estimator typically exceeds the observed richness by orders of magnitude (Fig. 3B). Notably, although multiple estimates of the naive TCR repertoire richness in a single individual can be quite heterogeneous, the estimated TCR richness is clearly distinct between both age groups. In addition, the rarefaction curves are mostly flat at the highest sampling depth, suggesting that the richness estimate is quite robust. We also integrated the information from multiple subsamples using the Chao2 estimator (Fig. 3C). The Chao2 estimates are based on occurrence in one or multiple samples rather than on the abundance in an individual sample [2]. The estimates did not saturate completely at the current sampling depth but were clearly distinct between the age groups, in line with the previous analyses [9]. In comparison with the Chao1 estimates, we arrived at much higher estimates for the total repertoire richness using the Chao2 estimator. This may indicate that the inferred cell abundance in individual samples is far from perfect, casting doubt on the accuracy of the Chao1 estimates in Fig. 3B. The total richness predicted using the Chao2 estimator appeared surprisingly consistent when based on a subset of the subsamples at a given sampling depth (Fig. 3D). So, the estimated richness based on multiple subsamples allows to discriminate between the naive TCR repertoires of young and aged individuals, even at a very limited coverage of the entire TCR diversity.

### Towards a robust comparison of diversity between sampled TCR repertoires

Here we analyzed the observed and extrapolated richness of multiple existing TCR repertoire sequencing datasets. While identifying ‘holes in the repertoire’ using TCR sequencing may be an attractive idea, the extremely limited coverage of typical samples weakens the outcomes of such an analysis. While estimating the richness of specific T-cell responses is problematic, even estimating the total richness of a diverse T-cell repertoire appears far from straightforward. Differences in richness and evenness may reflect differences in subset frequency rather than true differences in richness within a given T-cell pool. Addressing the changes in diversity during healthy ageing requires sequencing the TCR repertoire of multiple samples from sorted T-cell populations. It remains to be determined to which extent the observed or estimated TCR diversity can functionally explain or predict clinical outcomes in health and disease. This report illustrates that careful experiments and analyses are necessary to obtain robust signals from sampled immune repertoires.

## Methods

### TCRβ sequencing data

The processed TCRβ repertoires that were used for the analysis presented in Fig. 1 and Fig. 2, originally published in [6], were downloaded from the Adaptive Biotechnologies website. We only used the repertoires of donors with a known age and CMV-infection status, for which ‘counting method v2’ was applied. To eliminate the heterogeneity in sample size in Fig. 1, we used the 482 repertoires with a template count of at least 100 000, and down-sampled these without replacement to contain 100 000 templates. The TCR richness was quantified in these size-normalized repertoires, based on the combination of the identified V-gene, J-gene, and CDR3 nucleotide sequence. The dataset that was used for the analysis presented in Fig. 3, originally published in [9], was obtained from dbGaP found at https://www.ncbi.nlm.nih.gov/projects/gap/cgi-bin/study.cgi?studyid=phs000787.v1.p1 through dbGaP study accession number PRJNA258304. These data (project “Immunosenescence: Immunity in the Young and Aged”) were provided by Jorg Goronzy on behalf of his collaborators at PAVIR and Stanford University. In this study, five replicates with each 10^6^ cells per aliquot of naive and memory CD4 T cells were collected. For CD8 T cells, 0.25 *×* 10^6^ T cells were collected per replicate, except for the naive CD8 T cells from young individuals, from which 10^6^ cells per aliquot were used for sequencing. The sequencing data was processed using RTCR [7] as described before [5].

### Inference of a T-cell response against an unseen pathogen

The VDJdb [10] was downloaded from https://vdjdb.cdr3.net on 15 December 2022, by selecting human TRB sequences that were found to be specific for epitopes derived from HIV-1, with a confidence score of at least 1. When matching the size-normalized sequencing data with this database, we required the translated CDR3 sequence, as well as the V- and J-gene families to be identical.

### Reported naive T-cell counts

We searched the literature for studies reporting naive CD4 and CD8 T-cell counts as a function of age. We only selected studies in which individuals were stratified by CMV-status. We used the regression formulas presented in Table S1 in [12], the median ages and T-cell counts for both age groups reported in Tables 1 and 2 in [13], and the median T-cell counts plus standard deviation for the age groups reported in Tables 2 and 3 in [4].

**Table 1:** Example table showing observed and inferred TCRβ counts per subsample.

| TCRβ sequence | | | Observed number of reads | | | Inferred number of 'cells' | | |
| --- | --- | --- | --- | --- | --- | --- | --- | --- |
| V | J | CDR3 | Subsample 1 | ... | Subsample 5 | Subsample 1 | ... | Subsample 5 |
| 7-9 | 2-7 | TGTGCCA...GTACTTC | 0 | ... | 20 | 0 | ... | 2 |
| 7-9 | 2-3 | TGTGCCA...GTATTTT | 10 | ... | 7 | 1 | ... | 1 |
| 20-1 | 2-7 | TGCTGTA...GTACTTC | 0 | ... | 2 | 0 | ... | 1 |
| ... | ... | ... | ... | ... | ... | ... | ... | ... |
| 20-1 | 2-7 | TGCGGTG...GTACTTC | 1 | ... | 0 | 1 | ... | 0 |
| 9 | 1-1 | TGTGCCA...TTTCTTT | 38 | ... | 25 | 5 | ... | 3 |
| 7-9 | 2-3 | TGTGCCA...GTATTTT | 0 | ... | 20 | 0 | ... | 3 |

### Computational classification of total TCR repertoires into naive and effector/memory subpopulations

The dataset reported in [6] is based on unsorted T-cell repertoires. To address the potential changes in richness for the underlying T-cell subpopulations, we performed a TCR sequences by subset. We separated TCR sequences that are expected to be derived from naive T cells from those that were expected to be derived from effector/memory T cells. Specifically, we assumed the percentage of naive T cells in these samples to decrease from 90% at birth to 50% at 20 years of age (2 percent point decrease per year), and a further decrease of 0.25 percent point per year. We used the same selection criteria described above and inferred the total number of naive and effector/memory T cells in these samples. For the 420 TCR repertoires in which both numbers exceeded 100 000 cells (which in total thus contained at least 200 000 cells), we sorted the TCR repertoire by TCR sequence frequency. The least abundant sequences were classified as naive, until their relative frequency was equal to the estimated fraction of naive T cells. The remaining, more abundant, sequences were classified as being derived from effector/memory cells. Both fractions were down-sampled without replacement to a template count of 100 000 to eliminate the heterogeneity in sample size. The richness of each down-sampled TCR repertoire is shown in Fig. 2B.

### Inference of T-cell distributions from mRNA-based read distributions

The sequencing data obtained in [9] is based on PCR-amplified TCRβ mRNA transcripts from samples containing variable numbers of cells. The TCR frequencies in the data will thus be affected by differences in cell count, TCRβ expression level and amplification efficiency. These factors may influence the estimated richness in each of the samples, complicating reliable comparison between samples. To reduce the effect of these factors we inferred a distribution of cells from the distribution of reads in each sample. We reasoned that the majority of the TCR sequences observed in only one subsample will be derived from a single T cell. This allowed us to infer the distribution of how many reads are contributed by each single cell, for each subsample individually. Importantly, this distribution only describes the cells that contributed any TCR read, since cells without any contribution remain unnoticed. Using the average of the inferred number of reads that was derived from each contributing cell, we estimated how many cells contributed all reads together in each sample. Reassuringly, the subsamples containing naive CD8 T cells from aged individuals, that were known to contain fewer cells than the other samples, also contained fewer cells according to our TCR repertoire-driven estimate. We then assigned TCR sequences that were supported by a single read to single cells, since these reads cannot be contributed by multiple cells. The remaining TCR sequences were then assigned to inferred ‘cells’ based on the estimated read-contribution distribution, starting from the TCR sequence supported by the largest number of reads. This resulted in a table of TCR sequences, each supported by an inferred number of cells that was smaller than or equal to the observed number of reads supporting that TCR sequence (Table 1). The analyses presented in Fig. 3 are based on these inferred distributions of cells instead of the potentially more biased frequency of reads. Rarefaction curves were obtained by sampling without replacement from these inferred TCR repertoires.

## References

[1] Olga V. Britanova, Ekaterina V. Putintseva, Mikhail Shugay, Ekaterina M. Merzlyak, Maria A. Turchaninova, Dmitriy B. Staroverov, Dmitriy A. Bolotin, Sergey Lukyanov, Ekaterina A. Bogdanova, Ilgar Z. Mamedov, Yuri B. Lebedev, and Dmitriy M. Chudakov. Age-related decrease in TCR repertoire diversity measured with deep and normalized sequence profiling. The Journal of Immunology, 192:2689–2698, 2014.

[2] Anne Chao, Robert K. Colwell, Chih Wei Lin, and Nicholas J. Gotelli. Sufficient sampling for asymptotic minimum species richness estimators. Ecology, 90:1125–1133, 4 2009.

[3] Anne Chao, Yasuhiro Kubota, David Zeleńy, Chun Huo Chiu, Ching Feng Li, Buntarou Kusumoto, Moriaki Yasuhara, Simon Thorn, Chih Lin Wei, Mark J. Costello, and Robert K. Colwell. Quantifying sample completeness and comparing diversities among assemblages. Ecological Research, 35:292–314, 3 2020.

[4] S. Chidrawar, N. Khan, W. Wei, A. McLarnon, N. Smith, L. Nayak, and P. Moss. Cytomegalovirus-seropositivity has a profound influence on the magnitude of major lymphoid subsets within healthy individuals. Clinical and Experimental Immunology, 155:423–432, 2009.

[5] Peter C. de Greef and Rob J. de Boer. TCRβ rearrangements without a D segment are common, abundant, and public. Proceedings of the National Academy of Sciences, 118:e2104367118, 9 2021.

[6] Ryan O Emerson, William S DeWitt, Marissa Vignali, Jenna Gravley, Joyce K Hu, Edward J Osborne, Cindy Desmarais, Mark Klinger, Christopher S Carlson, John A Hansen, et al. Immunosequencing identifies signatures of cytomegalovirus exposure history and HLA-mediated effects on the T cell repertoire. Nature Genetics, 49:659, 2017.

[7] Bram Gerritsen, Aridaman Pandit, Arno C. Andeweg, and Rob J. de Boer. RTCR: a pipeline for complete and accurate recovery of T cell repertoires from high throughput sequencing data. Bioinformatics, 32(June):btw339, 2016.

[8] Chirag Krishna, Diego Chowell, Mithat Gönen, Yuval Elhanati, and Timothy A. Chan. Genetic and environmental determinants of human TCR repertoire diversity. Immunity and Ageing, 17:1–7, 9 2020.

[9] Qian Qi, Yi Liu, Yong Cheng, Jacob Glanville, David Zhang, Ji-Yeun Lee, Richard A. Olshen, Cornelia M. Weyand, Scott D. Boyd, and Jöorg J. Goronzy. Diversity and clonal selection in the human T-cell repertoire. Proceedings of the National Academy of Sciences, 111:13139–13144, 2014.

[10] Mikhail Shugay, Dmitriy V Bagaev, Ivan V Zvyagin, Renske M Vroomans, Jeremy Chase Crawford, Garry Dolton, Ekaterina A Komech, Anastasiya L Sycheva, Anna E Koneva, Evgeniy S Egorov, et al. Vdjdb: a curated database of T-cell receptor sequences with known antigen specificity. Nucleic acids research, 46:D419–D427, 2017.

[11] J. J. C. Thome, B. Grinshpun, B. V. Kumar, M. Kubota, Y. Ohmura, H. Lerner, G. D. Sempowski, Y. Shen, and D. L. Farber. Long-term maintenance of human naive T cells through in situ homeostasis in lymphoid tissue sites. Science Immunology, 1:eaah6506–eaah6506, 12 2016. DATA available via https://clients.adaptivebiotech.com/pub/2e838d8c-99cb-4103-b79e-7ee32768536d.

[12] A. M. Wertheimer, M. S. Bennett, B. Park, J. L. Uhrlaub, C. Martinez, V. Pulko, N. L. Currier, D. Nikolich-Zugich, J. Kaye, and J. Nikolich-Zugich. Aging and cytomegalovirus infection differentially and jointly affect distinct circulating T cell subsets in humans. The Journal of Immunology, 192:2143–2155, 2014.

[13] Liset Westera, Vera van Hoeven, Julia Drylewicz, Gerrit Spierenburg, Jeroen F. van Velzen, Rob J. de Boer, Kiki Tesselaar, and José A. M. Borghans. Lymphocyte maintenance during healthy aging requires no substantial alterations in cellular turnover. Aging Cell, 14:219–227, 2015.

[14] Eric J. Yager, Mushtaq Ahmed, Kathleen Lanzer, Troy D. Randall, David L. Woodland, and Marcia A. Blackman. Age-associated decline in T cell repertoire diversity leads to holes in the repertoire and impaired immunity to influenza virus. The Journal of Experimental Medicine, 205:711, 3 2008.

[15] Kengo Yoshida, John B. Cologne, Kismet Cordova, Munechika Misumi, Mika Yamaoka, Seishi Kyoizumi, Tomonori Hayashi, Harlan Robins, and Yoichiro Kusunoki. Aging-related changes in human T-cell repertoire over 20years delineated by deep sequencing of peripheral T-cell receptors. Experimental Gerontology, 96:29–37, 10 2017.

